# A hierarchical adaptive optics strategy for three-photon imaging during behavior

**DOI:** 10.1101/2025.05.04.652136

**Authors:** Huriye Atilgan, Jingyu Wang, Qi Hu, Sandra Tan, Blake Russell, Randy M. Bruno, Martin J. Booth, Armin Lak

## Abstract

Three-photon (3P) microscopy enables functional non-invasive single-cell resolution imaging at greater depths than any other technique. A key challenge of deep imaging is tissue-induced optical aberration, which reduces the excitation confinement. Adaptive optics use deformable mirrors to compensate for optical distortions, hence correcting these aberrations. Here, we present a practical adaptive optics-assisted 3P imaging system for functional imaging in the mouse brain during behavior. We introduce hierarchical corrections that sequentially target aberrations caused by the microscope system, the cranial window, and tissue depth. We demonstrate the utility of this strategy in the prelimbic cortex, where large vasculature near the midline causes aberrations, and in the lateral somatosensory cortex, where side access leads to distinct wavefront distortions. In both regions, adaptive optics significantly improved imaging performance, restoring cellular visibility near vasculature and enhancing signal-to-noise ratio. Our work provides a practical framework for utilizing adaptive optics to improve 3P imaging during behavior.

## Introduction

Recent advances in three-photon (3P) microscopy have demonstrated its potential for studying deep tissue neuronal activity underlying behavior (Horton et al., 2013; Ouzounov et al., 2017; Wang & Xu, 2020; Xiao et al., 2023). However, multi-photon imaging at depth presents substantial technical challenges due to the complex and heterogeneous optical environment of the brain. As light penetrates deeper into the tissue, it undergoes scattering and absorption, while also encountering wavefront distortions introduced by system misalignments and sample refractive index mismatches. These make it difficult to achieve a tightly focused laser beam for maximally efficient 3P excitation. Despite careful system alignment and sample preparation, small tilts of the cranial window, slight misplacements of optical elements (e.g., relay lenses, scanning mirrors, dichroic mirrors, and the objective), and inhomogeneities in the tissue can introduce wavefront distortions—or, in other words, phase aberrations. These aberrations degrade the point spread function and reduce the signal-to-noise ratio, ultimately limiting spatial resolution and image quality at depth.

Although 3P microscopy extends imaging depth due to reduced scattering and absorption at longer wavelengths (Takasaki et al., 2019), its performance is ultimately constrained by residual aberrations from both the microscope and the biological sample (Ji et al., 2012). System-induced aberrations, often arising from imperfect alignments within the microscope, can be corrected once using a static compensator. In contrast, sample-induced aberrations vary with imaging depth, brain region, and individual preparation, requiring dynamic and localized correction at the imaging plane. These aberrations, if uncorrected, lead to lateral blurring, axial elongation of the point spread function, and inevitably low signal strength. Therefore, precise wavefront correction is essential for maintaining image quality at depth. Adaptive optics offers a promising solution to these problems by actively correcting aberrations making it a critical tool for advanced 3P functional imaging (M. J. Booth, 2014; Hampson et al., 2021; Sinefeld et al., 2022; Streich et al., 2021; Xiaodong Tao et al., 2017).

Originally developed for astronomy, adaptive optics (AO) systems use deformable mirrors or other wavefront-shaping devices to compensate for optical distortions, allowing the excitation beam to maintain a tight focus in deep tissue. To assess and correct these distortions, both direct measurements of the wavefront and indirect image-based optimization methods can be used (Hampson et al., 2021). The latter approach, often preferred for its simplicity, modulates the phase of the laser beam by correcting common aberration patterns such as astigmatism, coma, and spherical aberrations from imaging metrics. By compensating for these distortions, adaptive optics significantly improves image sharpness, contrast, and signal strength, thereby enabling high-resolution functional imaging even in optically challenging heterogeneous tissues.

Here, we present a practical AO-assisted 3P imaging system optimized for in vivo functional recordings in the deep mouse cortex during behavior in head-fixed mice. To guide effective implementation of adaptive optics, we first introduce a hierarchical, three-level aberration correction strategy, targeting system-induced, cranial window-related, and deep tissue-specific distortions. We then demonstrate the necessity and application of this approach in two distinct imaging contexts: regions proximal to the superior sagittal sinus and lateral cortical regions. These cases illustrate how the AO system can be flexibly tuned depending on anatomical constraints and beam path geometry to achieve optimal imaging performance, showcasing the utility of adaptive optics in enhancing 3P imaging.

## Results

### Deep 3P imaging during behavior using adaptive optics

We begin by briefly describing our imaging system and the neuronal data we acquired in the frontal cortex during a Pavlovian conditioning task in head-fixed mice. We will then describe corrections that enhance the imaging quality before providing further examples to showcase the utility of our approach.

Our system utilized a Light Conversion Cronus-3P light source, integrating the pump laser, optical parametric amplifier, and prism compressor into a compact and efficient configuration. This system delivered a stable 1300 nm wavelength output, ideal for deep-tissue imaging applications. To further enhance image quality and correct optical aberrations, we incorporated a custom AO system. The beam was first expanded using a telescope before passing through a deformable mirror, with relay optics that conjugated a deformable mirror (DM) to the back pupil plane of the objective lens (**Figure 1A**; see Methods). Beyond the AO system, the imaging pipeline followed a conventional laser-scanning multiphoton microscope design: A pair of galvanometric scanners was used to acquire 256 × 256-pixel images at 4 Hz, which is necessarily limited by the lower repetition rates of 3P lasers. Fluorescence signals were collected with two photomultiplier tubes (PMTs) separated by a dichroic filter with a cut-off wavelength at 565 nm and coupled to a 16×, 0.8 NA Nikon objective lens (**Figure 1A**; see Methods), which provided a large numerical aperture for imaging depths of up to ~950 μm.

**Figure 1:**
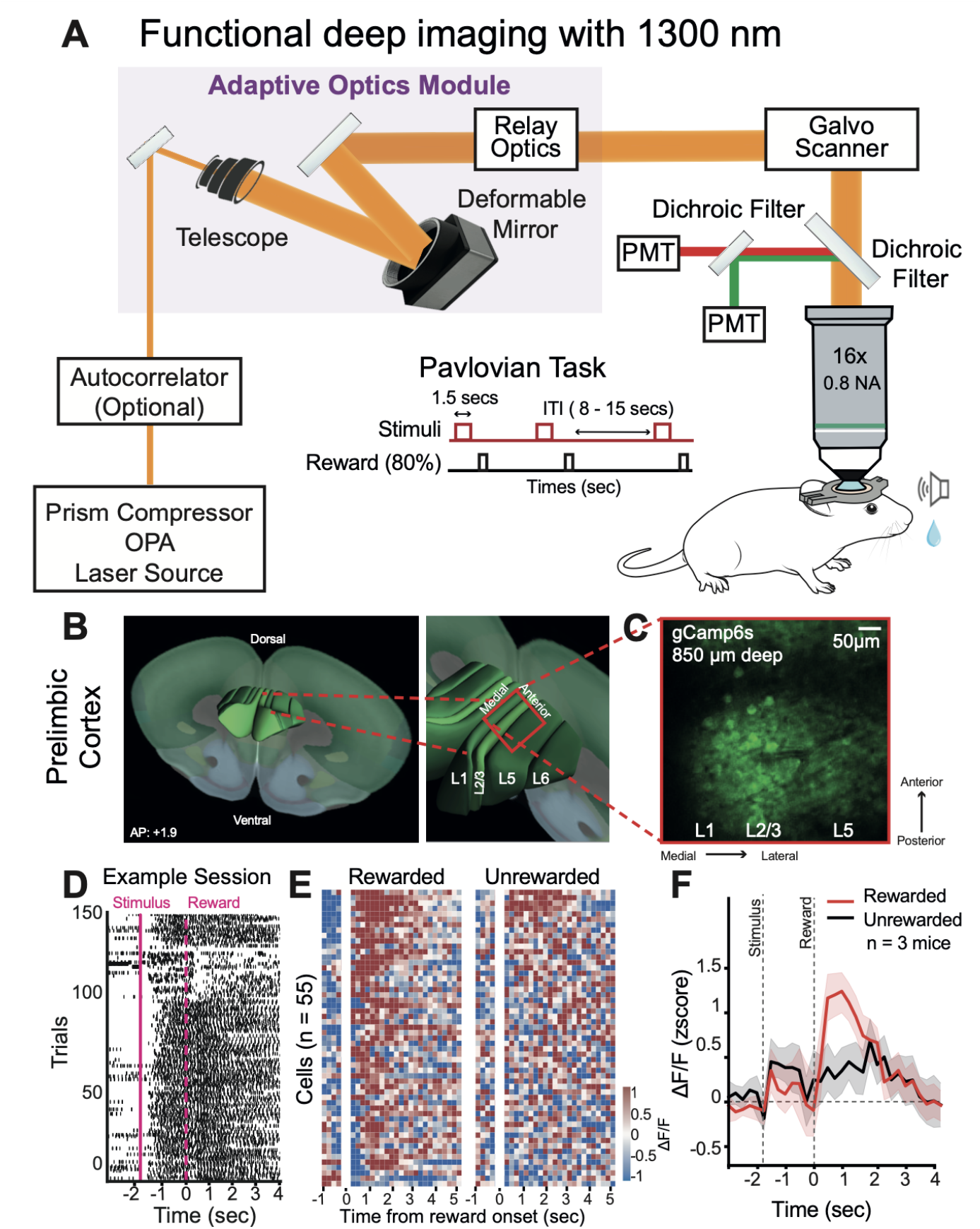
Functional deep imaging with 1300 nm wavelength illumination for in vivo recordings from the prelimbic cortex during behavioral performance in mice. **(A)** Schematic of the 3P imaging system, which includes an adaptive optics module for aberration correction, a telescope, a deformable mirror for beam shaping, and relay optics. The laser unit (Light Conversion CRONUS-3P) delivers 1300 nm light, which is directed through a galvo scanner and dichroic filters to two PMTs. A 16x, 0.8 NA Nikon objective lens is used for deep tissue imaging. **(B)** D model showing the placement of the cranial window over the prelimbic cortex in the mouse brain (AP = 141 +1.9), with a zoomed-in view highlighting cortical layers L1, L2/3, L5, and L6. **(C)** In vivo image from the prelimbic cortex at a depth of 850 µm, showing gCaMP6s expression in layers L1, L2/3, L5, and L6. **(D)** Example lick raster for an example session (n = 150 trials). Each row is one trial; dots mark individual licks (stimulus onset, solid magenta line at −1.5 seconds and reward delivery; dashed magenta line at 0 seconds). Anticipatory licks before reward reflect learned stimulus–reward associations. **(E)** Example session traces from rewarded (left) and unrewarded (right) trials, showing the change in fluorescence (ΔF/F) across cells (n = 55) recorded in the prelimbic cortex. The reward-aligned heatmap illustrates the temporal dynamics of neural activity during the auditory Pavlovian task. **(F)** Average neural responses to rewarded (red) and unrewarded (black) trials across 3 mice (n = 6 sessions, 2 sessions per mouse; 329 cells). Z-scored ΔF/F values are plotted over time, showing significant differences in neural activity during the stimulus and reward periods, demonstrating the system’s ability to capture decision-making processes in vivo.

To demonstrate the system’s performance, we imaged neuronal activity during behavior in head-fixed mice, focusing on the prelimbic cortex - a frontal cortical region that is implicated in reward valuation and decision-making but is not accessible using 2p imaging due to its depth (Palmer et al., 2024; Diehl & Redish, 2023; Lak et al., 2020; Sul et al., 2010). We first implanted a cranial window over the prelimbic cortex, located near the sagittal sinus and approximately 1.9 mm anterior to bregma. Imaging was conducted at 800–900 µm depths while mice performed a simple Pavlovian conditioning task. In this task, an auditory stimulus lasting 1.5 seconds was followed (0.5 seconds after the stimulus offset) by a liquid reward on 80% of trials (**Figure 1A**, see Methods). The scanning speed was set to 4 Hz due to the low repetition rate of the light source required for three-photon imaging; this rate was sufficient to capture task-related neural dynamics during the behavioral task.

We recorded the activity of several prelimbic neurons during the behavioral task. Despite cell-type-specific labeling of excitatory neurons with CamKIIα promoter-driven GCaMP6s expression, we observed notable somatic size variability within our field of view (FOVs, **Figure 1C**). This variability primarily arises because in the prelimbic areas, cortical layers are perpendicular to the imaging plane. Therefore, a single imaging plane encompasses neurons across multiple cortical layers (**Figure 1B and 1C**), each containing distinct pyramidal neuron subtypes and characteristic soma sizes (Benavides-Piccione et al., 2006; Gerfen et al., 2018; Harris & Shepherd, 2015; Molnár & Cheung, 2006). While optical aberrations at depth may also contribute to apparent neuronal size differences, the dominant source of variability in our recordings is the biological diversity across layers, spanning from superficial to deep layers within a single FOV. Given the substantial variation in cell size across cortical layers, we supplemented automatic cell detection using Suite2P (Pachitariu et al., 2016) with manual curation to ensure accurate identification of somatic masks (see Methods). Using this approach, we identified 329 cells across six sessions from three mice (mean = 55 cells/session, SD = 31; max = 119; min = 23).

To capture behavioral engagement, we recorded licking behavior during the task. Licks revealed tight temporal coupling to stimulus presentation, emerging prior to reward delivery and confirming that mice learned the stimulus–reward associations (**Figure 1D**). We then aligned neural activity to task events (i.e., stimuli and outcomes) and separated trials based on their outcomes (rewarded or unrewarded). Neuronal activity showed clear outcome-related divergence at the trial outcome, and also responded to reward-predicting cues in example sessions and across animals (**Figure 1E-F**). Our recordings uncovered rich neural dynamics during the task, suggesting that 3P imaging is a promising method for exploring how these dynamics vary across prelimbic layers in future studies. Overall, these results demonstrate that our AO-enhanced 3Photon system can reliably resolve task-related neural activity in deep cortical layers.

### Multi-level aberration compensation required for optimal imaging

Adaptive optics systems typically consist of a wavefront corrector—such as a deformable mirror (DM) — to modulate the wavefront, along with relay optics to direct the pre-compensated beam toward the sample. To determine the optical aberrations requiring correction, both direct wavefront sensing and indirect image-based optimization methods can be employed (Kristin N. Walker & Robert K. Tyson, 2009; Turaga & Holy, 2010; Wahl et al., 2019). The indirect - or sensorless - method of aberration measurement is particularly advantageous due to its simplicity and compatibility with standard fluorescence microscopy. It is often implemented using Zernike polynomials (Zernike, 1934), which represent common wavefront distortions such as coma, astigmatism, spherical, and trefoil aberrations. While higher-order aberrations also exist, these lower-order modes are the most amenable to correction with current DM technologies and are the primary contributors to image degradation at depth. The impact of these aberrations—and the improvement achieved by correcting them—is illustrated in Supplementary Figure 1.

In optimizing our system for deep imaging, we developed a hierarchical aberration correction protocol tailored for in vivo 3P imaging in the mouse brain (Table 1). This approach systematically rectifies aberrations introduced at different stages of the experimental setup, resulting in improved image quality and signal fidelity across imaging depths. 172

**Table 1:** Aberration correction protocol for deep tissue imaging.

| Aberration Source | Main aberration modes | Frequency | Protocol |
| --- | --- | --- | --- |
| 1. System-induced aberration | Coma, astigmatism, spherical | Once per imaging system | Use a fixed calibration slide. Position a cranial window used on the mice on a slide and get the best wavefront correction (called '*system flat*'). Apply these values to all recordings as default starting parameters. |
| 2. Head fixation-induced aberration | Coma, astigmatism | Once per animal | After applying the system flat, image a superficial layer of positioned mice and correct for residual coma and tilt introduced by head fixation. |
| 3. Deep tissue aberration | Trefoil, 2nd spherical, astigmatism, coma, and higher-order modes | Once per region of interest (ROI) | After correcting system and head fixation aberrations, locate your ROI. Run a few correction iterations to optimize the wavefront shape at depth. First, use large steps, then use the fine steps. |

#### 1. System-level aberration correction

We first addressed system-induced aberrations such as coma, astigmatism, and spherical aberrations. These aberrations are common in most microscopes: coma and astigmatism typically arise from optical misalignments, while spherical aberrations are primarily caused by the cranial window or the absence of a correction collar on the objective lens. These thus distort the point spread function and reduce imaging resolution and intensity (**Supplementary Figure 1**). To identify and correct these aberrations, we imaged a fixed slide containing 500 nm diameter yellow color fluorescent beads under a mounted cranial window (**Figure 2A**, top panel). Using this preparation, we optimized the adaptive optics system to generate a baseline correction, referred to as the “system flat.” This correction was applied uniformly across all animals imaged using the same optical setup and served as the foundation for all subsequent adjustments.

**Figure 2:**
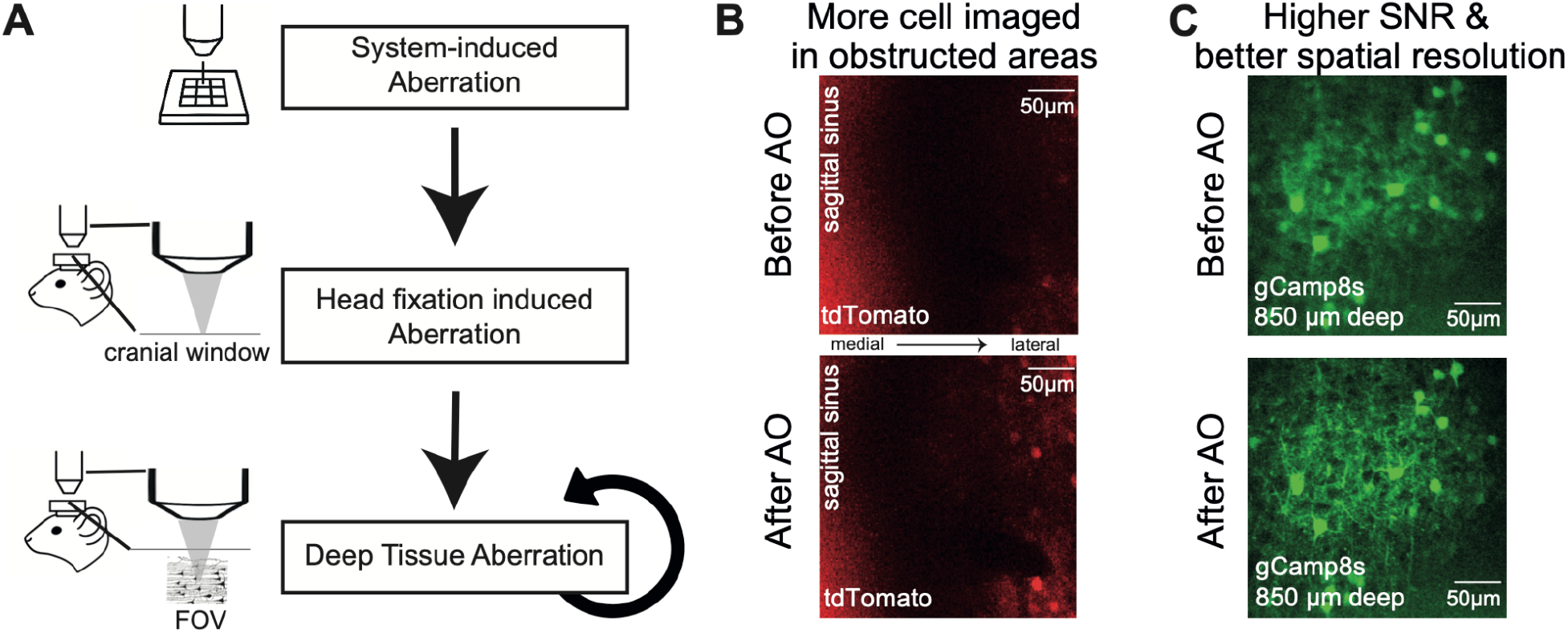
Multi-level aberration correction improves deep tissue imaging performance. **(A)** Schematic overview of the hierarchical aberration correction protocol. Aberrations were sequentially corrected at three stages: system-induced aberrations, head-fixation-induced aberrations, and deep tissue aberrations. **(B)** Representative in vivo tdTomato images near the sagittal sinus region before (top) and after (bottom) adaptive optics (AO) correction. Red emission detection under 1300 nm excitation enabled clear visualization of statically labeled cortical cells, improving image quality even in optically obstructed regions. AO correction led to a 36% increase in visible cells. **(C)** Representative in vivo GCaMP8s images at ~850 μm depth before (top) and after (bottom) full AO correction. AO correction substantially improved signal-to-noise ratio (15% increase) and enhanced spatial resolution.

The DM was pre-calibrated using a custom interferometer setup (Antonello et al., 2020; M. Booth et al., 2005) to obtain the control matrix for Zernike mode generation. To create the system flat, we corrected for coma, astigmatism, and spherical aberration. However, the exact pattern of aberrations depends on specific system misalignments. Therefore, we performed a comprehensive correction using Zernike modes — including both primary and secondary modes, up to mode 22. For each Zernike mode, we applied a sequence of known bias amplitudes (e.g., ±2 rad, ±1 rad, 0 rad), acquired images at each bias, and computed an image-quality metric based on fluorescence intensity. We then fitted a parabola to the measured intensity values to determine the optimal correction for each mode. This iterative sensorless AO routine, while time-consuming (requiring five measurements per mode, see Methods for details), was performed only once to generate a ‘system flat’ correction. The resulting correction was then saved and reused across imaging sessions.

#### 2. Head-fixation-induced aberration correction

Next, we corrected for aberrations introduced by slight variations in cranial window alignment during head fixation. Head-fixation errors typically result in minor tilts and thickness errors of the cranial window, which introduce aberrations such as coma and spherical aberrations. To correct for this, after applying the system flat, we imaged a superficial cortical plane to correct any remaining coma and spherical (**Figure 2A**, middle panel). To minimize preparation time and reduce the duration of animal handling during head fixation, we limited this correction step to three Zernike modes: primary spherical and two comas (*x* coma, *y* coma), with five aberration bias values tested per mode. This allowed efficient correction of superficial aberrations before deeper imaging. Head-fixation correction, performed once per animal, took around two minutes and prevented variability in head fixation from affecting imaging deeper regions.

Aberrations induced by head fixation could be corrected using both fluorescence signals of static labelling (e.g., red channel) and autofluorescence of surface cortical fibres captured during functional signals (e.g., green channel). Static labeling of neurons, such as with tdTomato, provides a stable signal that is inherently more reliable for intensity-based AO correction than the dynamic signals produced by calcium indicators like GCaMP, enabling more accurate aberration measurements. Nevertheless, in preparations only using dynamic GCaMP6s labeling, surface cortical fibers still exhibited strong autofluorescence, providing sufficient signal for effective correction even without additional static labeling (**Figure 3B**, left; an example for GCaMP6s cortical surface). Importantly, the DM is achromatic; therefore, applying the correction in one channel also improved image quality in the other, as a sharpened illumination focal spot improved fluorescence generation in all color channels equally.

**Figure 3:**
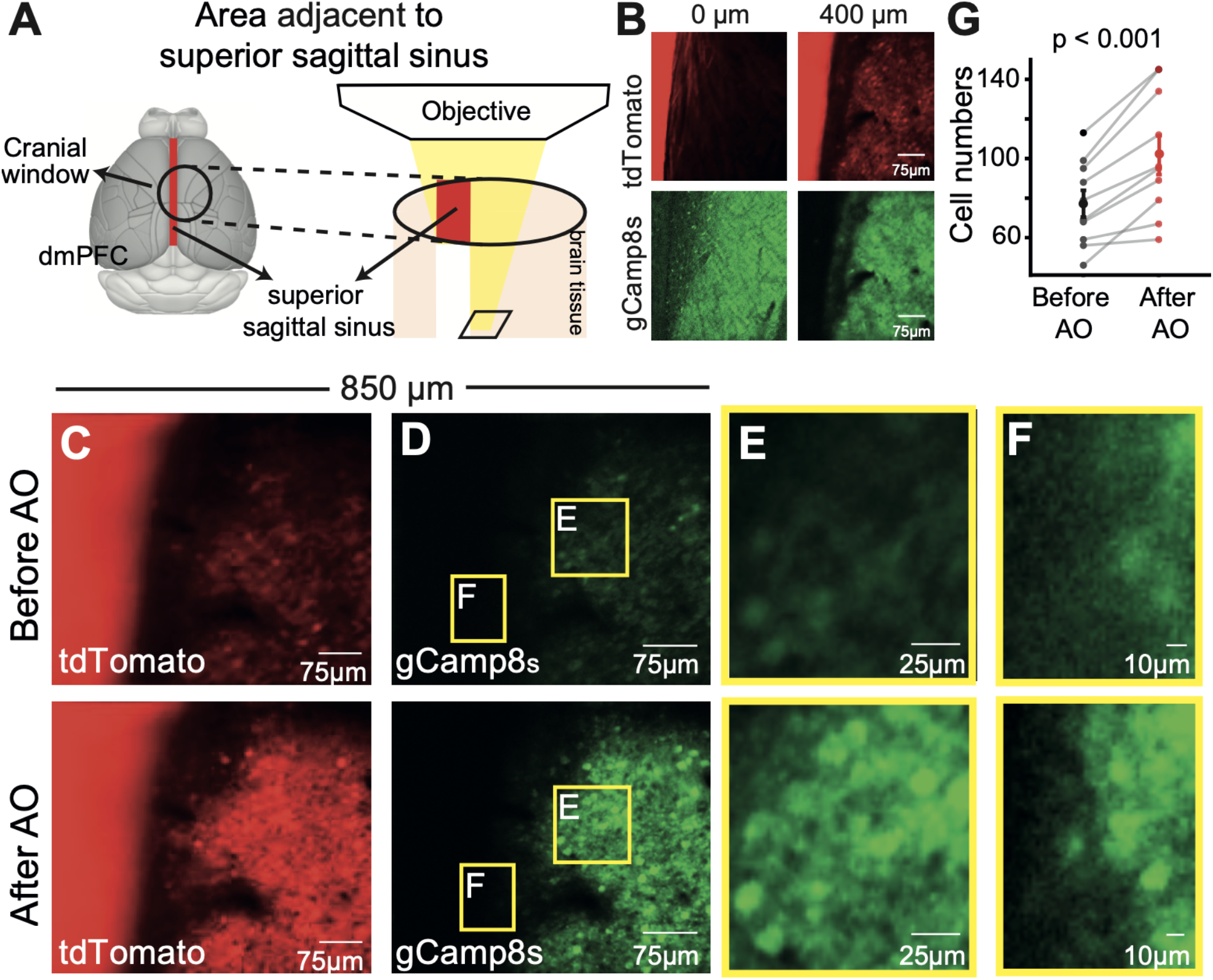
Adaptive optics improves cellular visibility and signal quality near the superior sagittal sinus. **(A)**Schematic showing imaging fields positioned adjacent to the superior sagittal sinus in the anterior cingulate cortex ACC) through a 4 mm cranial window. **(B)** tdTomato (top) and GCaMP8s (bottom) fluorescence images at the cortical surface (0 μm) and at 400 μm depth, illustrating the optical challenges imposed by the sinus region. **(C–D)** Representative tdTomato (C) and GCaMP8s **(D)** images at ~850 μm depth before (top) and after (bottom) adaptive optics (AO) correction. The same contrast settings were applied to the before and after images to allow direct visual comparison of signal improvement. **(E–F)** Magnified views of example subregions (E and F, indicated in yellow boxes in D) showing finer somatic structures before and after AO correction. Their contrast was optimised for each image. **(G)** Quantification of detected cell numbers before and after AO correction from three different mice (each data point represents one field of view). Cell detection was performed using Suite2p output combined with manual curation. Rigid motion correction (Suite2p) was applied to all recordings prior to analysis. Imaging was performed at 256 × 256 pixel resolution, 4 Hz frame rate, with 1-minute recordings averaged over 240 frames. Statistical comparison by paired t-test, p < 0.001.

#### 3. Deep tissue aberration correction

Finally, we addressed the complex and heterogeneous aberrations caused by refractive index variations due to inhomogeneous tissue structures in the deep brain. For this step, we corrected a total of seven aberration modes: four common lower-order aberrations (two coma and two astigmatism), trefoils, and secondary spherical aberrations. Following the first two correction steps, we targeted the region of interest (ROI) and applied iterative, image-based adaptive optics optimization to achieve precise, localized correction (**Figure 2A**, bottom panel). Although correcting additional higher-order aberrations could further improve image quality in deep cortical regions, doing so significantly increases optimization time. As such, to balance imaging quality and experimental practicality, we focused on correcting only seven key aberrations in our deep tissue protocol. A single correction run typically takes around 2 minutes; however, in more demanding imaging conditions, multiple iterations may be required to achieve optimal image quality, potentially extending the process to several tens of minutes. Nonetheless, depending on factors such as imaging depth, target structure, or optical configuration, correcting additional aberrations beyond our core set may further enhance imaging performance in other experimental setups.

Correcting the aberrations caused by tissue depth was best achieved using static imaging because many sensorless adaptive optics methods for multi-photon imaging require optimizing an image quality metric, which is often related to the fluorescence intensity (Rodríguez et al., 2021; Streich et al., 2021). Fluctuating fluorescence signals, such as the dynamic signals varying with neural activities, can affect the quality metric and make it unsuitable for adaptive optics optimization. Notably, there are image sharpness-based quality metrics that focus on measuring the image high frequency components. Such metrics, although not directly measuring the fluorescence intensity, can still be affected by varying fluorescence signals. GCaMP signals are dynamic and fluctuate with neural activity, and thus are not well-suited for estimating wavefront aberrations during AO optimization. Therefore, we co-expressed a static red fluorescent marker (tdTomato) within the ROI to guide sensorless adaptive optics correction. The presence of a stable, non-activity-dependent signal was critical for accurately measuring and refining the wavefront during iterative optimization (**Figure 3C and 3D**). Unlike GCaMP6s, whose signal varies with spontaneous and evoked activity, the static red fluorescence provided a consistent and robust output. This strategy substantially improved the accuracy and efficiency of deep tissue adaptive optics correction and enabled reliable restoration of high-resolution imaging at depth. Since both channels were excited by the same laser and the adaptive optics modulates the excitation path, correcting aberrations improved the overall focal quality, enhancing the signal in both channels.

Overall, this three-tier correction protocol enhanced cell visibility and improved imaging contrast, signal-to-noise ratio (SNR), and spatial resolution. In the representative example shown in **Figure 2B**, the number of visible cells increased from 14 to 34 (a 71% increase), and in **Figure 2C**, the SNR improved by 15%. Notably, the additional cells only became detectable following correction because their fluorescence intensity increased from sub-threshold levels to a visible level. These examples illustrate the qualitative and quantitative benefits of our approach in single FOVs. To assess these improvements systematically, we analyzed multiple imaging sites across animals, as detailed in **Figure 3** (for the case in **Figure 2B**) and **Figure 4** (for the case in **Figure 2C**), which report the distribution of improvements in cell counts, SNR, and resolution across fields of view and animals. By isolating and compensating for aberrations at each source—the optical system, head fixation, and brain tissue—we demonstrate that a sequential, targeted approach to aberration correction results in robust imaging performance.

**Figure 4:**
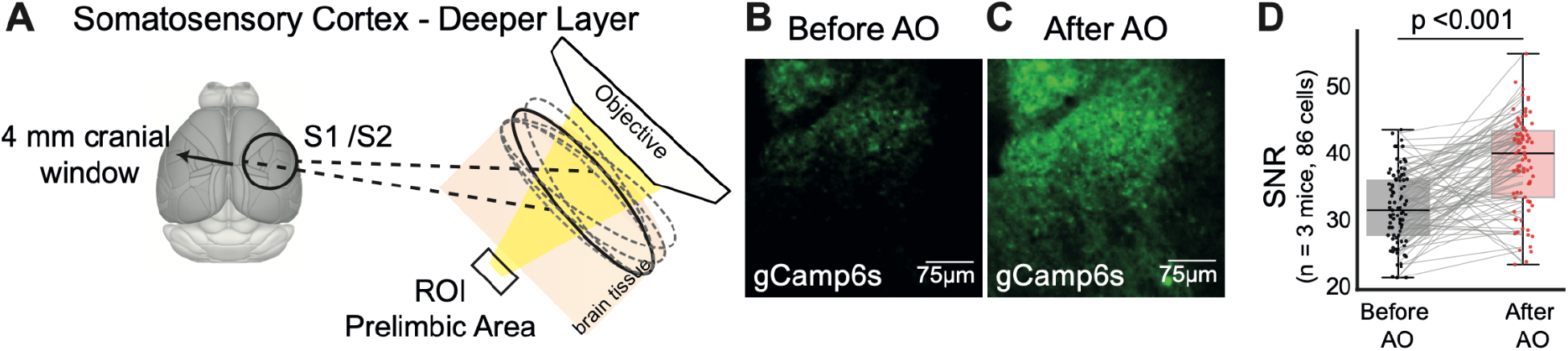
Adaptive optics improves signal-to-noise ratio during deep imaging in the lateral somatosensory cortex. **(A)** Schematic showing the imaging strategy targeting deep layers (~800–900 μm) of the lateral somatosensory cortex (S2) through a 4 mm cranial window. **(B–C)** Representative GCaMP6s images at ~850 μm depth before (B) and after (C) adaptive optics (AO) correction. The same contrast settings were applied to both images to allow direct comparison of signal enhancement following correction. **(D)** Quantification of signal-to-noise ratio (SNR) improvements for individual cells before and after AO correction. Each dot represents one cell (n = 86 cells, Suite2p ROI with manual curation); SNR was calculated from mean fluorescence over time divided by standard deviation across frames for each ROI. Rigid motion correction (Suite2p) was applied to all recordings prior to analysis. Imaging was performed at 256 × 256 pixel resolution, 4 Hz frame rate, with 1-minute recordings averaged over 240 frames. The vast majority of cells (≈85%) showed increased SNR, while some exhibited decreases due to the single AO correction applied to the whole FOV.

### Application of Aberration Correction Across Distinct Anatomical Contexts

To further evaluate the robustness of our correction strategies, we applied the protocol in two cortical regions that introduce distinct imaging challenges: the medial prelimbic cortex where its proximity to superior sagittal sinus causes difficulty in imaging **(Figure 3)** and the secondary somatosensory cortex, where imaging could be challenging due to lateral position of FOV (**Figure 4**).

In the medial prelimbic cortex, imaging fields were positioned close to the midline, where the superior sagittal sinus creates both anatomical and optical challenges at the cortical surface (**Figure 3A**). The sinus, a large blood-filled vessel, introduces significant refractive index heterogeneity and physical obstruction along the excitation path, complicating efforts to image deeper cortical regions (e.g., ~800 μm below the surface). At 1300 nm, some excitation light can partially transmit through the sinus, but this often results in substantial beam distortion, scattering, and signal loss. Additionally, the physical presence of the sinus displaces and deforms nearby cortical tissue, introducing local curvature and reducing the effective numerical aperture (NA) of the beam, further complicating access to otherwise viable ROIs. Depth-resolved imaging before correction revealed pronounced signal degradation and image distortion beginning as early as 300–500 μm depth (**Figure 3B**). Following full aberration correction at 850 μm depth (**Figure 3C-D**), we observed marked improvements in cell visibility (**Figure 3E**), even in areas adjacent to the sinus (**Figure 3F**).

Adaptive optics compensated for the wavefront distortions introduced by the complex optical environment near the superior sagittal sinus. This region presents a particularly challenging imaging context due to refractive index mismatches not only from the cranial window, but also from brain tissue (n ≈ 1.36–1.37) and blood (n ≈ 1.35–1.39, depending on oxygenation and hematocrit). These mismatches (especially when combined) cause localized phase delays, wavefront aberrations, and focal curvature across the imaging plane, leading to rapid degradation of spatial resolution and contrast, even at relatively shallow depths. By correcting low-order aberrations. (e.g., coma, astigmatism, spherical aberration, trefoil) based on the unique optical features of each imaging FOV, AO correction enabled finer somatic structures to re-emerge, significantly enhancing cell visibility even in regions adjacent to major vasculature. Although the tissue absorption cannot be recovered, the benefits of AO correction were consistently replicable across different ROIs within and across mice, as quantified by improved signal fidelity in 10 ROIs from 3 mice (**Figure 3G;** paired t-test, t_19_= 3.567, p < 332 0.001).

We next applied our multi-level aberration correction strategy to recordings in the lateral somatosensory cortex, a representative example of more lateral cortical areas (**Figure 4A**). In contrast to imaging near the midline, the lateral cortex presents a more uniform surface morphology, with relatively fewer large surface vessels and minimal gross tissue curvature. However, imaging at depths of 800–900 μm still exhibited noticeable degradation in spatial resolution and signal intensity prior to aberration correction (**Figure 4B**). These degradations were primarily driven by deep tissue scattering and slight cranial window tilts introduced during surgical preparation, where the window was not perfectly perpendicular to the optical axis of the objective, as is often unavoidable.

Application of the full correction protocol significantly enhanced imaging quality, as evidenced by improved signal-to-noise ratio (**Figure 4D**, pairwise t-test t_85_ = −7.65; *p* < 0.001) While the total number of visually detectable somata remained similar before and after correction (**Figure 4B-C**), AO correction substantially improved the clarity, contrast, and reliability of individual cell signals. The vast majority of cells (~85%) showed increased SNR, while some exhibited decreases due to the single AO correction applied to the whole FOV (see Discussion for sub-region tuning within FOV methods). This suggests that, even in lateral cortical regions without major superficial vascular structures, multi-level aberration correction is essential for maximizing functional imaging fidelity at depth.

## Discussion

Our results demonstrate that integrating adaptive optics with 3P microscopy significantly enhances deep brain functional imaging capabilities. By systematically correcting optical aberrations of the system, cranial window, and tissue, we were able to extend functional imaging depths while preserving high spatiotemporal resolution. This simplified correction strategy allowed us to reliably record neuronal activity during stimulus-guided behavior from brain regions where fluorescent imaging was previously not possible or severely degraded by optical distortions. In developing these corrections, we paid special attention to experimental practicality and time constraints, ensuring that the corrections can be applied in the context of experiments typically run on mice to study the relationship between neuronal signals and behavior.

While alternative strategies such as microprisms and gradient-index (GRIN) lenses have enabled access to deep brain structures (Andermann et al., 2013; Barretto et al., 2009), they require invasive implantation and may distort tissue anatomy or limit the usable field of view. In contrast, 3P microscopy allows for volumetric access to deep cortical and subcortical areas with minimal tissue disruption. Its longer excitation wavelengths and reduced scattering make it particularly well-suited for functional imaging in intact brain tissue, especially when combined with adaptive optics to overcome remaining aberrations.

We show that the anatomical context strongly influences the distribution of aberrations encountered during deep imaging. In the medial prelimbic cortex, close to the superior sagittal sinus, large surface blood vessels introduce strong absorption and wavefront distortions through refractive index heterogeneity, leading to rapid signal degradation even at shallow depths. In contrast, in the lateral somatosensory cortex, although superficial morphology was more regular, subtle cranial window tilts and cumulative deep tissue still substantially compromised image quality at depth. AO correction markedly improved imaging performance in both contexts: restoring cellular visibility adjacent to vasculature in the prelimbic cortex and enhancing signal-to-noise ratios in the lateral cortex, even though the number of detectable cells remained constant in the latter.

Our data also highlight an important biological and geometric consideration: in regions such as the medial prefrontal cortex, deep cortical layers exhibit greater apparent heterogeneity in cell size, reflecting both diverse pyramidal subtypes and the curvature of the cortical sheet at depth. Because automated ROI detection algorithms can be biased by soma size, this variability poses a potential confound for both detection efficiency and downstream population-level analyses. Even in the face of such heterogeneity, AO remains critical by restoring diffraction-limited resolution. AO helps to delineate true cell boundaries across varied soma sizes. Recognizing and accounting for cell-size heterogeneity will therefore be essential in future 3P functional imaging studies of deep cortical circuits, ensuring that observed activity patterns truly reflect neural dynamics rather than detection artifacts.

We demonstrated that correcting low-order Zernike aberrations (coma, astigmatism, spherical aberration, trefoil) already delivers substantial gains in image quality and scientific insight for deep 3P imaging. Looking ahead, extending this framework to higher-order modes— especially with deformable mirrors or spatial light modulators offering greater actuator density—could further counteract scattering-induced distortions and unlock even finer resolution at depth. Likewise, adopting automated optimization strategies, such as machine-learning–based AO (Hu et al., 2023), would accelerate aberration correction and dynamically adapt to complex wavefront errors. Although we chose a conventional sensorless approach to preserve simplicity and retain precise control over individual aberration components, our protocol is fully compatible with these advanced methods — paving the way for more efficient, robust, and high-fidelity deep-tissue imaging in future studies. Traditional wavefront sensing has been successfully used in two-photon imaging (Liu et al., 2019), but it becomes challenging at great depths because of the reduced photon budgets in 3P imaging.

Like most adaptive optics microscopy approaches, here we applied a single correction pattern across each field of view—a choice that delivers clear benefits in simplicity and overall image quality, as demonstrated by our deep-tissue results. When optical aberrations vary substantially within a large FOV, for example, around small blood vessels or local scattering centers, region-specific corrections may yield even higher fidelity. Future implementations could synchronize the DM with the scanners for sub-region tuning, or incorporate multi-conjugate AO and direct wavefront-sensing techniques to achieve field-dependent aberration correction. By moving toward spatially adaptive AO strategies (Hampson et al., 2020; Rajaeipour et al., 2020; Wu & Cui, 2015; Mertz et al., 2015; Simmonds & Booth, 2013; Kam et al., 2007), it will be possible to preserve the advantages of our current framework while achieving optimal aberration compensation across large, heterogeneous brain regions.

Overall, our findings emphasize that adaptive optics is a transformative tool for deep in vivo imaging during systems neuroscience experiments, enabling high-quality functional recordings across anatomically diverse and optically challenging brain regions. With further hardware and software innovations, particularly around expanded correction modes and smarter depth-adaptive control, AO-assisted 3P microscopy holds substantial potential to push the boundaries of functional neuroimaging deeper into the intact brain.

## Methods

### Animals

All experimental procedures involving animals were conducted in accordance with the UK Animals in Scientific Procedures Act (1986). Adult mice of both sexes (2-6 months of age) on a C57BL/6 background were used for experiments. We used seven transgenic mice on a CamKIIα–tTa (AI94) × B6; DBA-Tg(tetO-GCaMP) background—two expressing GCaMP8s and five expressing GCaMP6s. All mice were housed at room temperature (20–22 °C) on a standard light/dark cycle and humidity of ~40%, and provided with a standard mouse chow diet and received daily husbandry.

### Surgical Procedures

Three different surgical procedures were used in this project: (1) implantation of a cranial window over the sinus, (2) implantation of a temporal window over the sinus combined with a retrograde viral injection into the dorsomedial striatum (DMS) to label projection neurons in the cortex, and (3) implantation of a lateral temporal window over the right somatosensory cortex.

All animals underwent the same anesthetic protocol. Anesthesia was induced with 3% isoflurane and maintained at or below 1.5% throughout surgery. Perioperative analgesia included subcutaneous administration of buprenorphine (0.1 mg/kg, Vetergesic) and meloxicam (5 mg/kg, Metacam). Additionally, bupivacaine (2 mg/kg, Marcaine) was applied topically to the scalp prior to sterilization with chlorhexidine gluconate and isopropyl alcohol (ChloraPrep). The scalp was then bilaterally removed. The exposed skull was cleaned with a bone scraper (Fine Science Tools) to remove the periosteum. An aluminum head-plate with a 7 mm imaging well was secured to the skull using dental cement (Super-Bond C&B, Sun-Medical).

Depending on the experimental condition, mice underwent a circular craniotomy (4 mm diameter over right somatosensory cortex: –1.4 mm posterior, +4.0 mm lateral; or 4-5 mm diameter over the sagittal sinus targeting prelimbic cortex: +1.5 mm anterior, centered over the sinus) performed with a dental drill (NSK UK Ltd.). The skull within the craniotomy site was soaked with saline before removal. Bleeding, if present, was flushed with saline for at least 5 minutes before performing a durotomy.

In animals undergoing viral labelling, a single 500 nL viral injection was performed using a calibrated, sharpened glass pipette. To label DMS-projecting neurons in the prelimbic cortex, a retrograde virus expressing red fluorescent protein (rAAV2-CAG-tdTomato, 2.3× 10^13^ gc/mL, 500 nL, Addgene #59462-AAVrg) was injected into the DMS (AP +0.74, ML +1.25, DV 1.8). This injection strategy was originally designed to enable deep imaging of the striatum (DV = 1.8) and to push the limits of our imaging protocol. A useful consequence of this approach is that it produces sparse labeling across different cortical depths as well as in the striatum, which proved advantageous for testing the effectiveness of our aberration correction method. Injections were delivered at a rate of 100 nL/min using an automated nanoinjector (Nanoject II™, Drummond). Following the injection, the pipette was left in place for 10 minutes to promote viral diffusion.

After the injection, a double-layered cranial window—composed of a 4 mm coverslip glued to a 3 mm coverslip (or a 5 mm glued to a 4 mm coverslip)—was gently pressed into the craniotomy and sealed with cyanoacrylate (VetBond) and dental cement. Mice were placed in a heated recovery chamber and monitored until fully ambulatory. Postoperative care included daily health monitoring and weight recording for 7 days.

Mice were allowed a minimum of 21 days recovery with ad libitum access to food and water prior to any further procedures, allowing sufficient time for viral expression before the onset of behavioral training.

### Behavioral training

Pavlovian conditioning was employed as the training task to familiarize the auditory cue to the mice, utilizing a 1.5-second tone followed by a water reward delivered in 80% of the trials. Over a 3-5 day training period, animals were subjected to 150 trials per day, with each trial separated by an inter-trial interval (ITI) ranging from 8 to 15 seconds. 49

### Three-photon imaging

The three-photon fluorescence microscope utilized an integrated Cronus-3P Unit (Light Conversion, Photonics Solutions, UK), which combines a wavelength-tunable optical parametric amplifier (OPA) and a prism compressor, pumped by a femtosecond laser (1030 nm). The system operated at 1300 nm and achieved a pulse width of 49 fs under the objective lens. The laser repetition rate was maintained at 1 MHz with a maximum power above 100 mW under the objective lens. The images were acquired with a galvo scanning multiphoton microscope (VivoScope, Scientifica UK) and a water immersion objective (Nikon, 16x, 0.8 NA, 3mm working distance). We also tested a higher numerical aperture, long working distance objective (Olympus XLPLN25XSVMP2, 25×, 1.0 NA, 4 mm working distance), but observed no substantial improvement in imaging quality at these depths. Although larger NA objectives offer better transmission, theoretical benefits in resolution and collection efficiency, they are physically bulkier, making precise positioning over the cranial window more challenging. Achieving an optimal working distance often requires a larger cranial window, which in turn introduces greater susceptibility to motion artifacts. As a result, the practical complications associated with using a larger NA objective outweighed the potential optical advantages in our deep imaging experiments.

One PMT (H10770PA-40, Hamamatsu) detection channel was used to collect green fluorescence using a dichroic (T565LPXR, Chroma) and bandpass filter (ET525/50m-2p, Chroma). The second PMT detection channel was used to collect red fluorescence with a bandpass filter (ET620/60m, Chroma). The signal was then converted to voltage by a transimpedance amplifier (150 kOhm, 1 MHz; XPG-ADC-PREAMP) and sent to a channel of the digitizer of the vDAQ card (Vidrio Technologies). ScanImage 2021 (Vidrio Technologies) running on MATLAB (MathWorks) was used to acquire images. A motorized stage (Scientifica) was used to control the position of the sample, and a focusing motor (Scientifica) was used to control the position of the objective and PMT detection apparatus relative to the sample. All imaging depths are reported in raw axial movement of the motorized stage that holds the objective lens. High-resolution structural images were typically taken with 256×256 pixels per frame at 3.2 μs dwell time (4.23 Hz) and an average of 240 frames unless otherwise stated.

### Adaptive Optics

An adaptive-optics Mirao 52e deformable mirror (DM) was incorporated into the excitation path. The laser beam after the OPA was first expanded to fill the DM aperture, then relayed by a telescope to conjugate the DM to the scanners and the back pupil plane of the objective lens. This pupil conjugation guaranteed that wavefront corrections were applied uniformly across the entire field of view in our telecentric design. To match the beam diameter to the DM’s 15 mm aperture and the 3mm scanning mirrors, the expanded laser beam was reflected from the DM and passed through an 8.3x (500/60) telescope. The beam diameter was reduced to 1.8 mm (defined at the 10% power cut-off) at the scanners. After the scanners, the beam was expanded by a telescope formed by the scan lens and tube lens to a diameter of 12.75 mm at the objective pupil plane. This underfilled the objective’s back aperture (16.67 mm diameter), corresponding to a fill ratio of 0.765 for a Gaussian beam, calculated based on the 10% power cut-off definition.

Before first use, the continuous-membrane DM must undergo a thorough Zernike polynomial control matrix calibration to account for individual actuator influence function. We performed this calibration on a homebuilt interferometer (design and code available at https://aomicroscopy.org/dm-calib, (Antonello et al., 2020)), where each actuator was “poked” through a range of voltages while capturing the resulting surface profiles. Custom software (https://github.com/dop-oxford/dmlib) then extracted wavefronts from the interferograms and computed each actuator’s influence function. By fitting these influence functions to a set of normalized Zernike modes, we assembled and inverted a conversion matrix that maps desired Zernike polynomials to actuator voltages. This automated pipeline ensured that any prescribed wavefront correction—expressed in Zernike terms—could be delivered precisely by the DM.

During experiments, we implemented a sensorless adaptive-optics routine to correct residual aberrations without adding extra hardware or special sample preparations. Sensorless adaptive optics treats image quality metric as the feedback signal: because optimal aberration correction maximizes image intensity, sharpness, or contrast, we define an image metric— such as the fluorescence intensity or the normalized variance or high-frequency power of the fluorescence image—and then iteratively apply an intentionally biased Zernike mode on the DM to measure that metric. For each mode, we applied a set of Zernike biases, acquired an image, computed its metric value, and used a quadratic function to fit the measured curve to converge on the optimal correction. This flexible, purely image-based approach scales seamlessly across depths and sample types; for full implementation details, see Zhang et al. (2023).

In this work, we adopted a modal sensorless adaptive optics strategy, which is well-suited to our continuous‐membrane DM and practical for routine use. We chose the three‐photon fluorescence intensity as our image‐quality metric, since improving the point‐spread function directly increases three‐photon yield. To ensure that metric changes reflected aberration correction rather than FOV shifts, we excluded the tip, tilt, and defocus Zernike modes—each of which translates the FOV laterally or axially—from our optimization. For each remaining aberration mode, we applied a series of known bias amplitudes and acquired images over the same fixed FOV, then computed the three‐photon intensity for each bias. In the small‐ aberration regime (<1 rad root mean square), the metric’s dependence on mode amplitude is approximately parabolic, so we fitted a parabola to these data points and located the peak bias as the optimal correction. Because Zernike modes are orthogonal over a circular pupil, assuming with even illumination, correcting one mode does not, in theory, alter the others, enabling a straightforward sequential optimization. Variations on this approach—and a discussion of its advantages and limitations—can be found in Facomprez et al. (2012).

In this paper, we used the 5N correction scheme (meaning five different known biases per mode were introduced for correction derivation). We selected two comas, two astigmatisms, two trefoils, and primary spherical as modes for correction. For each mode, the five biases are ±2 rad, ±1 rad, and 0 rad to allow an equally spaced coverage over the total range of 2 rad, which is adopted in early 2P imaging studies Facomprez et al., (2012). Other smart sensorless AO approaches have also been demonstrated recently to be successful in correcting aberrations and even partially for scattering in three-photon imaging at great depths (Berlage et al., 2021; Qin et al., 2022).

### Data Analysis

All data analyses were performed using custom scripts in MATLAB and Python unless otherwise noted. Image processing was intentionally kept minimal to preserve the integrity of the raw data. No spatial smoothing, sharpening, or deconvolution was applied. Images shown in figures primarily represent the average intensity projection of raw or minimally processed frames, aiming to accurately reflect native imaging quality. Motion correction was applied using Suite2p’s non-rigid motion correction algorithm (Pachitariu et al., 2016), allowing for the correction of both global and local movements across imaging frames. This step was critical to maintaining structural consistency, particularly during deep tissue imaging.

Cell numbers were determined by manually curated detection of somatic masks, initially generated using Suite2p’s automated segmentation. Given the variability in soma size across cortical layers, automatic detection alone was insufficient. Automated segmentation outputs were manually reviewed and adjusted to ensure accurate identification of cells. Therefore, we combined Suite2p output with manual curation to ensure accurate identification of somatic cell masks. The total number of identified cells was then quantified for each field of view and used 600 in comparative analyses. Statistical comparisons of cell count across conditions were also 601 performed using two-tailed paired t-tests. Signal-to-noise ratio (SNR) was calculated by 602 dividing the mean fluorescence intensity of the signal region, estimated from Suite2p-identified 603 somatic masks with neuropil subtraction applied, by the standard deviation of the background 604 signal within the same field of view. SNR comparisons across conditions were statistically 605 evaluated using two-tailed paired t-tests.

To quantify the improvement in fluorescence signal following AO correction, we measured 608 total pixel intensity in raw images acquired before and after AO correction. Images were first 609 converted to 16-bit grayscale using Fiji (ImageJ). Total intensity was quantified as the 610 Integrated Density (sum of pixel values within the field) via the “Analyze > Measure” function. 611 The percentage change in intensity was calculated by comparing the values obtained from 612 AO-before and AO-after images.

### Data and software availability

The functional three-photon calcium imaging recordings that are presented in this manuscript 615 will be made publicly available in a GIN repository upon publication. Behavioral training used 616 Rigbox hardware and software as previously reported (Bhagat et al., 2020). Offline pre-617 processing imaging analysis was performed using FIJI (Schindelin et al., 2012) and Suite2p. 618 All subsequent data analysis and visualization were performed in Python 3.9 using custom-619 written code.

## Acknowledgments

We are grateful to Light Conversion, Photonics Solutions (UK), for kindly lending us the 623 CRONUS-3P laser unit. This work is supported by a Sir Henry Wellcome postdoc fellowship 624 (222807/Z/21/Z) to H. A., grants from Wellcome (213465/Z/18/Z) and UKRI (EP/X026655/1) 625 to A.L., UKRI/Wellcome Physics of Life Grant (EP/W024047/1), European Research Council 626 Advanced Grant (AdOMiS 695140) to M.J.B., Schmidt Sciences LLC (Schmidt AI in Science 627 postdoctoral research fellowship) to Q.H., and a Wellcome Trust Discovery Award 628 (225210/Z/22/Z) to R.M.B.

## Figures & Legend

**Supplementary Figure 1:**
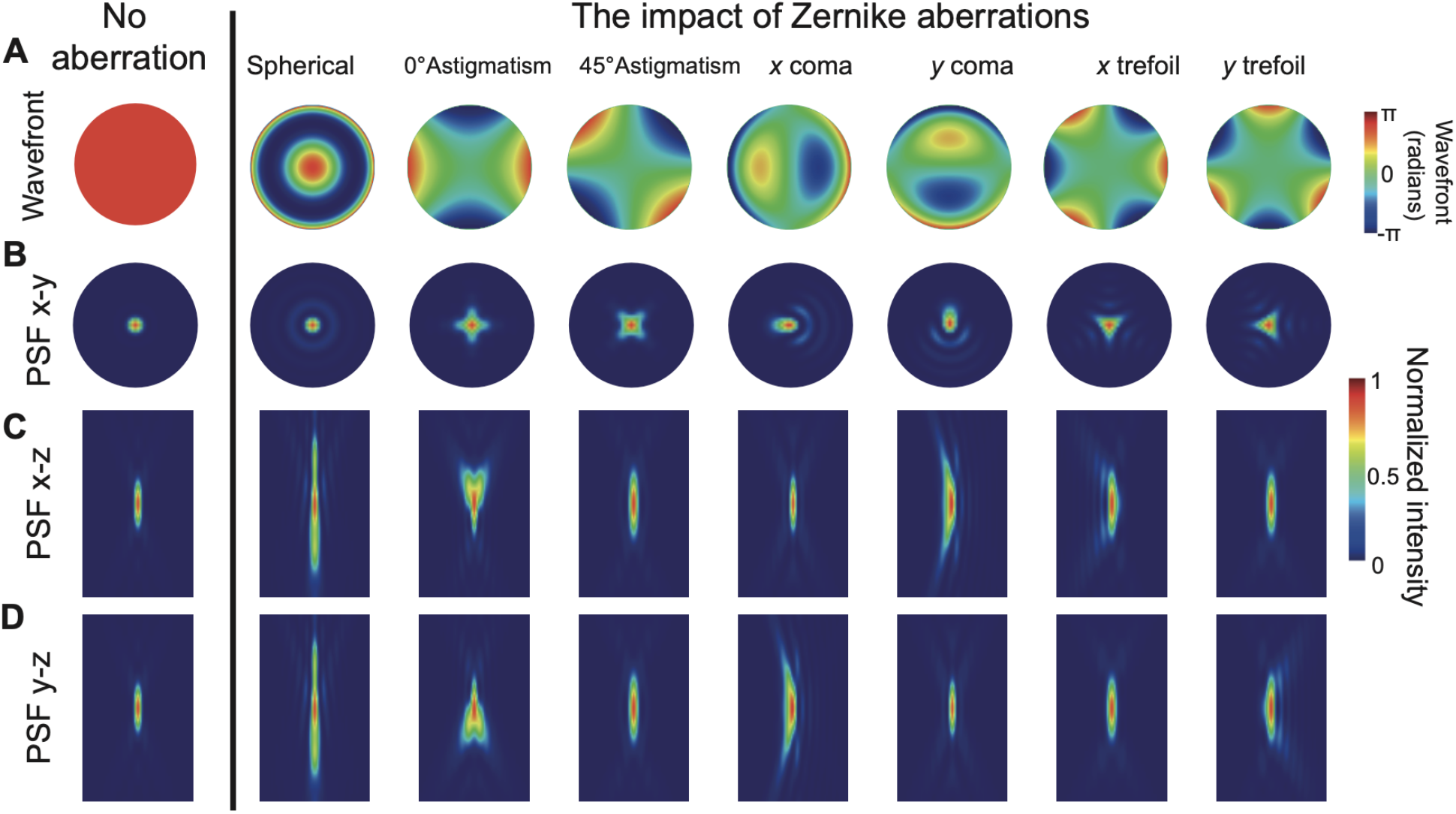
Impact of individual Zernike aberrations on the wavefront and point spread function (PSF). **(A)** Simulated wavefronts for a set of low-order Zernike modes, including: piston (no 634 aberration), spherical aberration, 0° and 45° astigmatism, *x* and *y* coma, and *x* and *y* trefoil. 635 The piston mode represents a uniform wavefront with no distortion. **(B–D)** Corresponding 636 simulated PSFs showing the effect of each aberration on image quality in three orthogonal 637 planes: **(B)** lateral plane (x–y), **(C)** axial x–z plane, and **(D)** axial y–z plane. Each aberration 638 produces characteristic PSF distortions, demonstrating its impact on spatial resolution and 639 optical performance. The aberrations generated were all at 1 radiance root-mean-square 640 (RMS) value. Scalebar for wavefront maps (row A) represents 1 numerical aperture (NA) unit 641 across the pupil plane. Scalebars for PSF images (rows B–D) represent 1 µm in the lateral (x– 642 y) views and 2 µm in the axial (x–z and y–z) views.

## Notes

### Competing Interest Statement

The authors have declared no competing interest.

### Summary of Updates

All text is updated to clarify the adaptive protocol; Figures 2, 3, and 4 revised; more analytical results added; author affiliations updated; Supplemental figure added.

